# The Spike Proteins of SARS-CoV-2 B.1.617 and B.1.618 Variants Identified in India Provide Partial Resistance to Vaccine-elicited and Therapeutic Monoclonal Antibodies

**DOI:** 10.1101/2021.05.14.444076

**Authors:** Takuya Tada, Hao Zhou, Belinda M. Dcosta, Marie I. Samanovic, Mark J. Mulligan, Nathaniel R. Landau

## Abstract

Highly transmissible SARS-CoV-2 variants recently identified in India designated B.1.617 and B.1.618 have mutations within the spike protein that may contribute to their increased transmissibility and that could potentially result in re-infection or resistance to vaccine-elicited antibody. B.1.617 encodes a spike protein with mutations L452R, E484Q, D614G and P681R while the B.1.618 spike has mutations Δ145-146, E484K and D614G. We generated lentiviruses pseudotyped by the variant proteins and determined their resistance to neutralization by convalescent sera, vaccine-elicited antibodies and therapeutic monoclonal antibodies. Viruses with B.1.617 and B.1.618 spike were neutralized with a 2-5-fold decrease in titer by convalescent sera and vaccine-elicited antibodies. The E484Q and E484K versions were neutralized with a 2-4-fold decrease in titer. Virus with the B.1.617 spike protein was neutralized with a 4.7-fold decrease in titer by the Regeneron monoclonal antibody cocktail as a result of the L452R mutation. The modest neutralization resistance of the variant spike proteins to vaccine elicited antibody suggests that current vaccines will remain protective against the B.1.617 and B.1.618 variants.

## Introduction

Despite efforts to contain severe acute respiratory syndrome coronavirus-2 (SARS-CoV-2) the virus has continued to spread throughout the world’s population, generating novel variants with mutations selected for immunoevasion and increased transmissibility. The variants contain point mutations and short deletions in the spike protein driven by selective pressure for increased affinity for its receptor, ACE2, and escape from neutralizing antibody. Major variants identified to date include B.1.1.7 [1, 2], B.1.351 [3], B.1.526 [4], P.1 [5], P.3 (EpiCoVTM database of the Global Initiative for Sharing All Influenza Data (GISAID) : accession numbers EPI_ISL_1122426 to EPI_ISL_ 1122458) and mink cluster 5 [6]. The variants have increased transmissibility, a feature that is, at least in part, the result of mutations in the spike protein that allow for increased ACE2 affinity and/or resistance to antibody. Several of the point mutations have been found to lead to increased viral fitness. The N501Y mutation in B.1.1.7 results in increased affinity for ACE2 [7-10] while the E484K mutation in the B.1.351, B.1.526, P.1 and P.3 spike proteins provides partial resistance to neutralizing antibodies in recovered individuals and antibodies elicited by vaccination [11-16].

Recent months have seen a dramatic increase in India in the rate of spread of SARS-CoV-2 infection accompanied by an increase in mortality. The increased spread is associated with newly identified novel SARS-CoV-2 variants B.1.617 [17] and B.1.618 (http://cov-lineages.org) with mutated spike proteins. The B.1.617 spike protein has L452R, E484Q, D614G and P681R mutations [17]; and the B.1.618 spike has mutations Δ145-146, E484K and D614G. Residues 452 and 484 are in the receptor binding domain (RBD) and thus could play a role in immunoevasion and/or resistance to antibody neutralization. Δ145-146 lies in the N-terminal domain that is a known antibody binding site [18] and P681R lies within S1/S2 where it could affect proteolytic processing **(Fig. 1A and B)**. Which of the mutations contributes to increased ACE2 affinity and resistance to antibody neutralization that might account for the increased transmissibility of the variants is not well understood.

**Figure 1.**
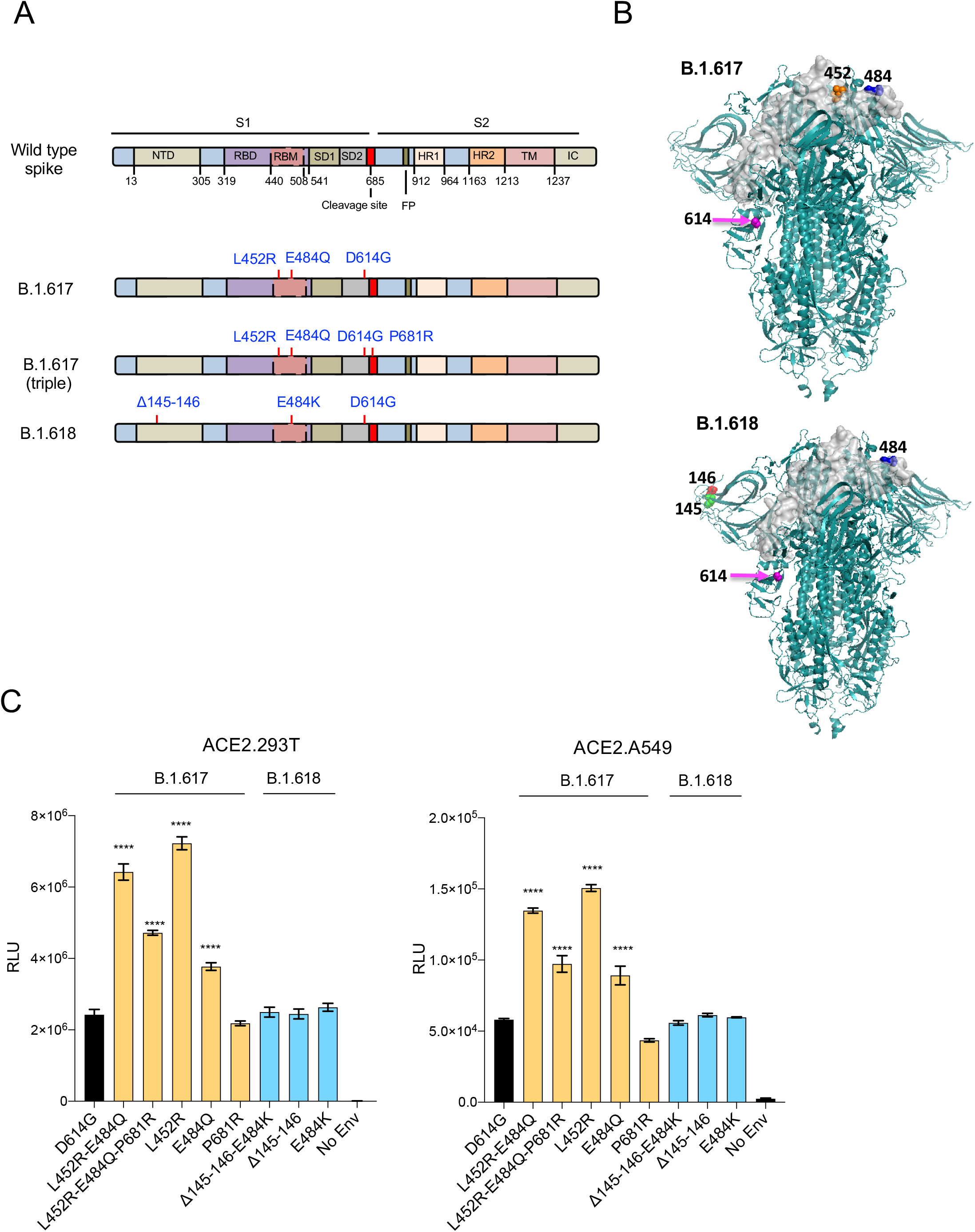
Infectivity of virus pseudotyped by B.1.617 and B.1.618 variant spikes. (A) The diagram shows the domain structure of the SARS-CoV-2 spikes of B.1.617 and B.1.618. NTD, N-terminal domain; RBD, receptor-binding domain; RBM, receptor-binding motif; SD1 subdomain 1; SD2, subdomain 2; FP, fusion peptide; HR1, heptad repeat 1; HR2, heptad repeat 2; TM, transmembrane region; IC, intracellular domain. (B) The 3D images indicate the location of the key mutations in B.1.617 and B.1.618 spikes. One representative RBD is shown in gray for simplicity. (C) Expression vectors for the variant spike proteins, deleted for the carboxy-terminal 19 amino acids, were generated and used to produce pseudotyped viruses. Viruses were normalized for RT activity and applied to ACE2.293T cells. Infectivity of viruses pseudotyped with the individual mutations of the B.1.617 and B.1.618 spike protein or combinations thereof were tested on ACE2.293T (left) and ACE2.A549 cells (right). Luciferase activity was measured two days post-infection.

In this study, we addressed the questions of antibody resistance and variant spike protein affinity for ACE2 using lentiviruses pseudotyped by the B.1.617 and B.1.618 spike proteins. We found that the viruses with the Indian spike proteins were partially resistant to neutralization by convalescent serum antibody and vaccine-elicited antibodies. The resistance was caused by the L452R, E484Q and E484K mutations. In addition, the variants were partially resistant to REGN10933, one of the two mAbs constituting the Regeneron COV2 therapy.

## Results

### Generation of B.1.617 and B.1.618 spike protein-pseudotyped lentiviruses

The B.1.617 variant spike protein contains L452R and E484Q mutations in the RBD in addition to D614G and the P681R mutation near the proteolytic processing site **(Fig. 1A and B)**. The B.1.618 spike has E484K in the RBD in addition to D614G and the N-terminal deletion Δ145-146 **(Fig. 1A and B)**. We constructed expression vectors for the B.1.617 and B.1.618 spike proteins and for spike proteins with the individual point mutations and used these to produce lentiviral pseudotypes that with a genome encoding GFP and luciferase reporters. Immunoblot analysis of transfected cells and supernatants showed that the variant spike proteins were expressed and proteolytically processed and that the spike proteins were incorporated into lentiviral virions at a level similar to that of wild-type D614G **(Supplementary Fig. 1)**. Quantification of the band intensities showed that the P681R mutation, which lies near the proteolytic processing site, caused a small increase in proteolytic processing as measured by a 2-fold decrease in the ratio of S/S2. Analysis of the infectivity of each virus, normalized for particle number, on ACE2.293T cells showed that the B.1.617 spike protein (L452R/E484Q/P681R) was >2-fold increase in infectivity while B.1.618 was similar to wild-type D614G. Analysis of the individual mutations showed that the increased infectivity of the B.1.617 spike was attributed to L452R, which itself caused a 3.5-fold increase in infectivity and in combination with E484Q caused a 3-fold increase. The other individual point mutations had no significant effect on infectivity (Δ145-146, E484K, P681R) (**Fig. 1C**). Analysis of the same panel of pseudotype viruses on ACE2.A549 cells showed a similar pattern of relative infectivity of each spike protein with an overall decrease of 50-fold infectivity.

### Neutralization of the Indian variants by convalescent sera and vaccine-elicited antibody

To determine the sensitivity of the Indian variants to antibody neutralization, we tested the serum specimens from convalescent patients who had been infected prior to the emergence of the variants for neutralization of the panel of pseudotyped viruses. The results showed that viruses with the B.1.617 and B.1.618 spikes were 2.3 and 2.5-fold resistant to neutralization by convalescent sera compared to wild type - a finding that was similar to that of the 3-fold resistance of the South Africa B.1.351 variant **(Fig. 2A)**. The resistance of B.1.617 was caused by the L452R and E484Q mutation and B.1.618 resistance was caused by the E484K mutation, as is the case for B.1.351. Δ145-146 and P681R had no significant effect on neutralization resistance.

**Figure 2.**
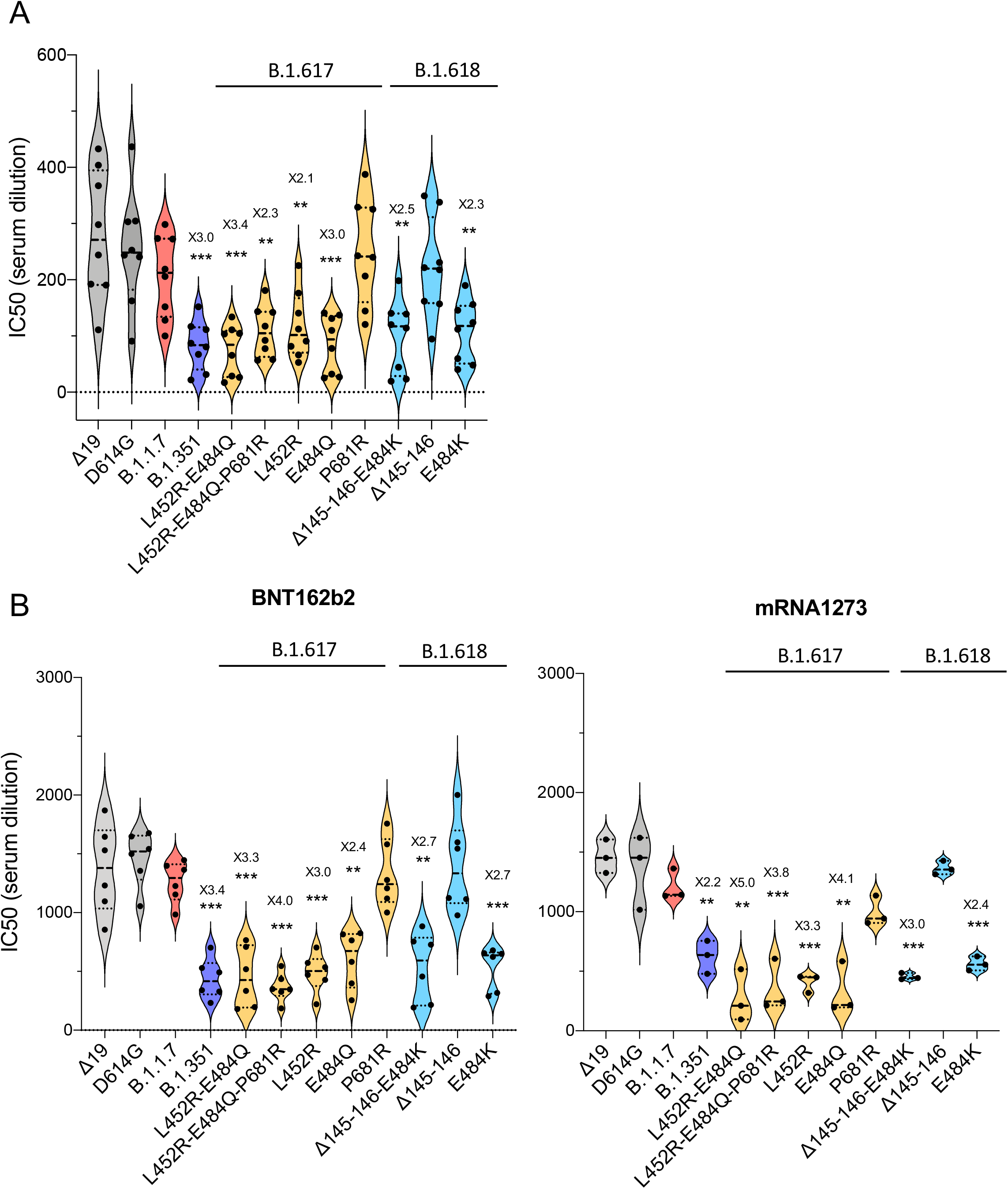
Neutralization of spike protein variants by convalescent sera and antibodies elicited by BNT162b2 and mRNA1273 vaccine. (A) Neutralization of viruses pseudotyped by B.1.617 (yellow) and B.1.618 (light blue) spikes (double, triple and single) by convalescent serum samples from 8 donors was tested. Each dot represents the IC50 for a single donor. The analyses were repeated twice with similar results. (B) Neutralizing titers of serum samples from BNT162b2 vaccinated individuals (n=6) (left) and mRNA1273 vaccinated donors (n=3) (right) was measured. IC50 of neutralization of virus from individual donors are shown.

To determine the resistance of the variants to neutralization by vaccine-elicited antibodies, we tested sera from individuals vaccinated with Pfizer BNT162b2 and Moderna mRNA-1273 vaccines for their ability to neutralize the panel of variant spike pseudotyped viruses. The results showed a similar pattern of resistance to neutralization as for the convalescent sera except that overall antibody titers were about 5-fold higher for sera from vaccinated individuals. B.1.617 was about 4-fold-resistant and B.1.618 was about 2.7-fold resistant to neutralization. The resistance could be attributed to L452R and E484Q in B.1.617 and E484K in B.1.618 **(Fig. 2B)**.

### Variant pseudotype neutralization by Regeneron REGN10933 and REGN10987 mAbs

Monoclonal antibody therapy for COVID-19 has been shown to reduce disease symptoms and to reduce the number of patients requiring hospitalization [19]. However, the treatment is subject to becoming less effective in patients infected with a variant in which the antibody epitope on the spike protein is altered by mutation. To address this question, we tested the ability of the mAbs to neutralize the panel of variant spike protein pseudotyped viruses. The results showed that the neutralizing titers of REGN10933 for B.1.617 virus was decreased by about 20-fold, similar to that of the E484K variant **(Fig. 3A, D)**. Δ145-146 and P681R did not affect neutralizing titer. Neutralizing titers of REGN10987 for viruses with the L452R and B.1.617 were decreased by about 3-fold (**Fig. 3B, D)**. The neutralizing titer of the mixture of REGN10933 and REGN10987 was 4.7-fold decreased in neutralizing titer for virus with the B.1.617 spike while the neutralization of virus with the B.1.618 spike was unchanged **(Fig. 3C, D).**

**Figure 3.**
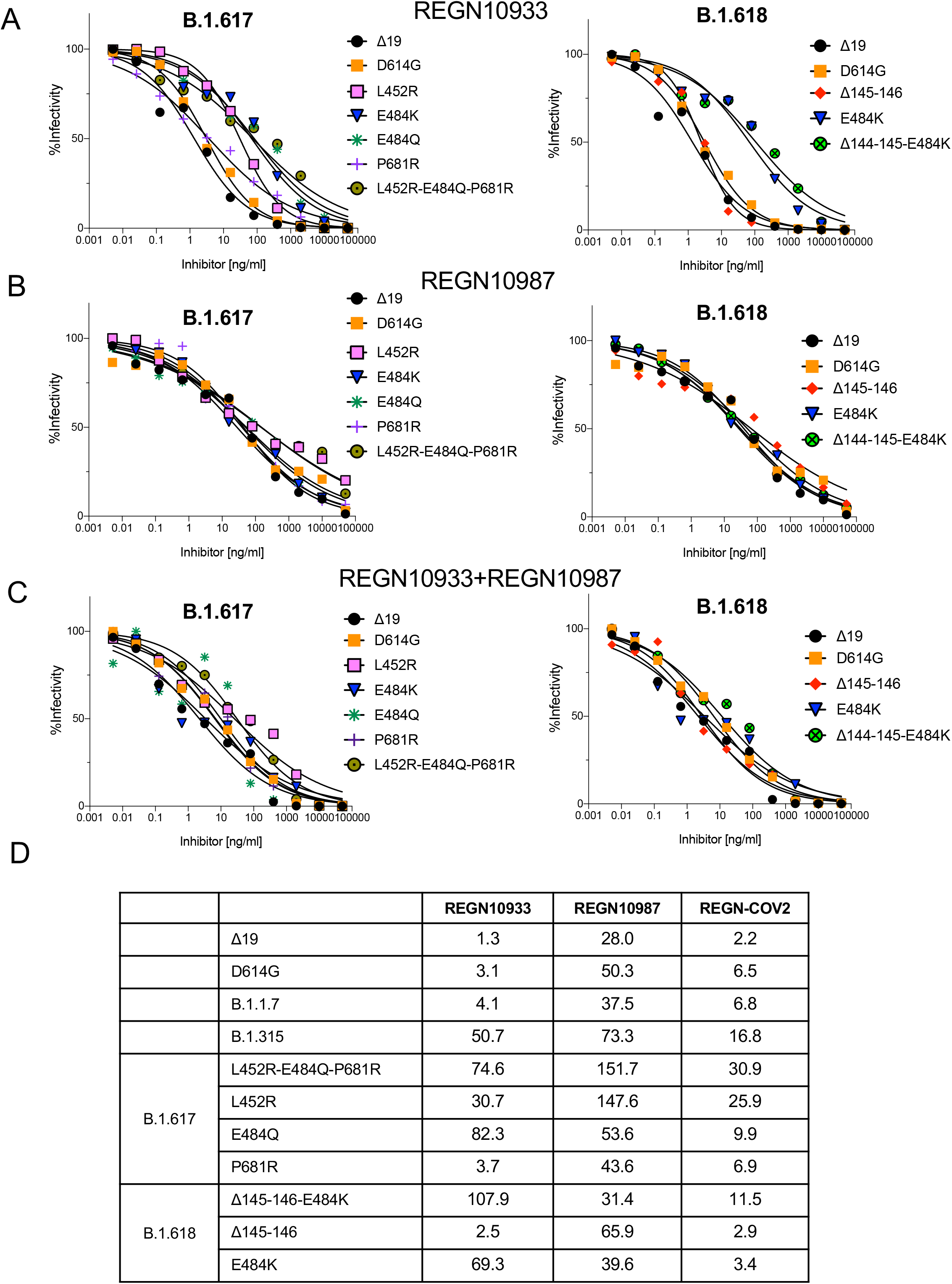
Neutralization of B.1.617 and B.1.618 spike protein variants by REGN10933 and REGN10987. Neutralization of B.1.617 (left) and B.1.618 (right) pseudotyped viruses by REGN10933 (A) and REGN10987 (B). (C) Neutralization of viruses pseudotyped by B.1.617 (left) and B.1.618 (right) spike proteins by 1:1 mixture of REGN10933 and REGN10987 was measured. (D) The IC50 from REGN10933, REGN10987 and combination antibodies was shown.

### B.1.617, B.1.618 have increased affinity for ACE2

The apparent increased transmissibility of the variants could be caused by increased affinity for ACE2 as has been found for other variants. To determine the relative ACE2 affinities of the variant spikes, we used an ACE2 binding assay in which the pseudotyped viruses were incubated with soluble ACE2 (sACE2) and then tested for infectivity on ACE2.293T cells relative to that of virus with the D614G mutation, an assay that we have used previously to analyze variant spike protein affinity [11]. The assay shows a 6-fold increase in ACE2 affinity for the N501Y mutation, a result consistent with that reported previously **(Fig. 4)**. The results showed only a small effect for the single E484K mutations but a larger effect for variants with the L452R or combination of E484Q and L452R. The P681R and Δ145-146 had no effect.

**Figure 4.**
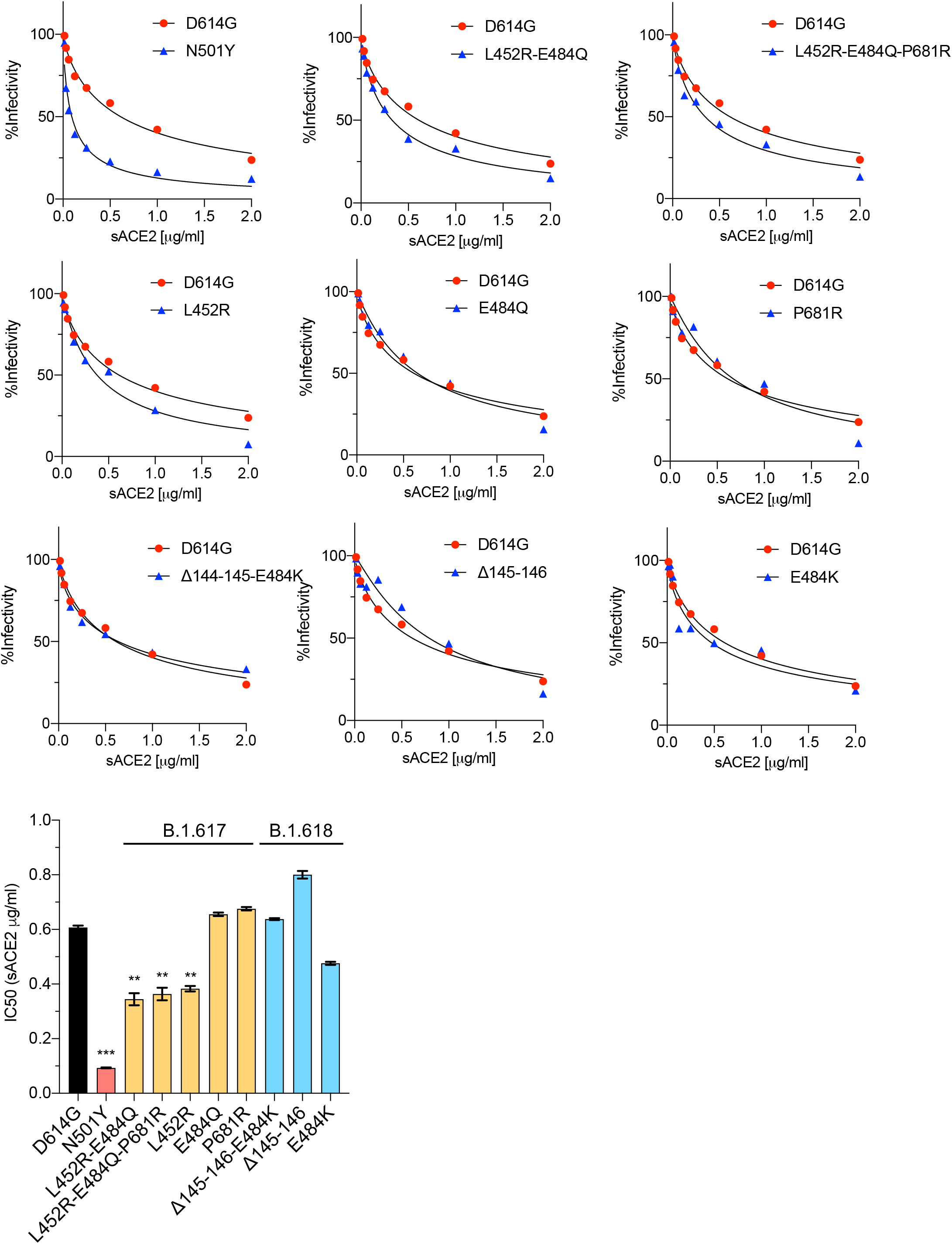
Neutralization of B.1.617 and B.1.618 spike protein variants by soluble ACE2. Viruses pseudotyped with variant spike proteins were incubated with a serially diluted recombinant soluble ACE2 (sACE2) and then applied to ACE2.293T cells. Each plot represents the percent infectivity of D614G (red) and B.1.617 and B.1.618 (blue) pseudotyped virus. The histogram at the bottom shows the calculated IC50 for each of the curves.

## Discussion

Viruses with the B.1.617 and B.1.618 spike were partially resistant to neutralization, with an average 3.9-fold and 2.7-fold decrease in IC50 for convalescent sera and antibodies elicited by Pfizer and Moderna mRNA vaccines, respectively. The neutralization resistance was mediated by the L452R, E484Q and E484K mutations. The extent of resistance of the variants was similar to that of the earlier B.1.351 and the New York B.1.526 (E484K) variant. The L452R and E484Q mutations provide an increased affinity for binding to ACE2, likely contributing to the increased transmissibility of the variants. Both variants were partially resistant to REGN10933, one of the two monoclonal antibodies constituting the REGN-COV2 therapy and virus with the B.1.617 spike was partially resistant to REGN10987 as well, resulting in a 4.7-fold decrease in neutralizing titer for the antibody cocktail. Even with the 3-4-fold decrease in neutralization titer of vaccine elicited antibodies, average titers were around 1:500, a titer well above that found in the sera of individuals who have recovered from infection with earlier unmutated viruses. Thus, there is a good reason to believe that vaccinated individuals will remain protected against the B.1.617 and B.1.618 variants.

The L452R mutation, which is present in the California B.1.427/B.1.429 was found to have a significant effect on resistance to vaccine-elicited and monoclonal antibodies. The variant was found to be shed with 2-fold increase in infected individuals, increase viral infectivity in cell cultures and confer a 4 to 6.7-fold and 2-fold decrease in neutralizing titers of antibodies from convalescent and vaccinated donors, respectively [20]. The E484K mutation present in the B.1.351, B.1.526, P.1 and P.3 spike proteins has been shown to cause partial resistance to neutralization as compared to the earlier D614G spike protein [11-16]. Mutation of the same position, E484Q, in B.1.617 caused a 2-4-fold decrease of neutralization by serum, demonstrating the importance of this residue as an epitope for antibody recognition. The P681R mutation at the furin cleavage site of B.1.617 is similar to the P681H mutation found in the B.1.1.7 spike proteins. P681R appeared caused a detectable increase in the amount of cleaved spike protein on virions. The increase was not associated with an increase in infectivity on ACE2.293T cells but might have an effect on the infection in primary cells *in vivo*.

The analyses in this study were restricted to the mRNA-based vaccines but there is no reason to believe that vector-based vaccines such as that of Johnson and Johnson that express a stabilized, native, full-length spike protein would be different with regarding to antibody neutralization of virus variants. Our results lend confidence that current vaccines will provide protection against variants identified to date. However, the results do not preclude the possibility that variants that are more resistant to current vaccines will emerge. The findings highlight the importance of wide-spread adoption of vaccination which will both protect individuals from disease, decrease virus spread and slow the emergence of novel variants.

## Methods

### Plasmids

B.1.617 and B.1.618 spike mutations were introduced into pcCOV2.Δ19.D614GS by overlap extension PCR and confirmed by DNA nucleotide sequencing. Plasmids used in the production of lentiviral pseudotypes have been previously described [22].

### Cells

293T cells were cultured in Dulbecco’s modified Eagle medium (DMEM) supplemented with 10% fetal bovine serum (FBS) and 1% penicillin/streptomycin (P/S) at 37°C in 5% CO_2_. ACE2.293T is a clonal cell-line that stably expresses a transfected human ACE2 as previously described. The cells were maintained in DMEM/1 μg/ml puromycin/10% FBS/1% P/S.

### Human Sera and monoclonal antibodies

Convalescent sera and sera from BNT162b2 or Moderna-vaccinated individuals were collected on day 28 following the second immunization at the NYU Vaccine Center with written consent under IRB approved protocols (IRB 18-02035 and IRB 18-02037). Donor age and gender were not reported. Regeneron monoclonal antibodies (REGN10933 and REGN10987) were prepared as previously described [21].

### SARS-CoV-2 spike lentiviral pseudotypes

Lentiviral pseudotypes with variant SARS-CoV-2 spikes were produced as previously reported [21]. Viruses were concentrated by ultracentrifugation and normalized for reverse transcriptase (RT) activity. To determine neutralizing antibody titers, sera or mAbs were serially diluted 2-fold and then incubated with pseudotyped virus (approximately 2.5 × 10^7^ cps) for 30 minutes at room temperature and then added to ACE2.293T cells. Luciferase activity was measured as previously described [22].

### Soluble ACE2 Neutralization assay

Serially diluted recombinant soluble ACE2 protein [22] was incubated with pseudotyped virus for 30 minutes at room temperature and added to 1 × 10^4^ ACE2.293T cells. After 2 days, luciferase activity was measured using Nano-Glo luciferase substrate (Nanolight) in an Envision 2103 microplate luminometer (PerkinElmer).

### Immunoblot analysis

Spike proteins were analyzed on immunoblots probed with anti-spike mAb (1A9) (GeneTex), anti-p24 mAb (AG3.0) and anti-GAPDH mAb (Life Technologies) followed by goat anti-mouse HRP-conjugated second antibody (Sigma) as previously described [22].

### Quantification and Statistical Analysis

All experiments were in technical duplicates or triplicates and the data were analyzed using GraphPad Prism 8. Statistical significance was determined by the two-tailed, unpaired t-test. Significance was based on two-sided testing and attributed to p< 0.05. Confidence intervals are shown as the mean ± SD or SEM. (*P≤0.05, **P≤0.01, ***P≤0.001, ****P≤0.0001). The PDB file of D614G SARS-CoV-2 spike protein (7BNM) was downloaded from the Protein Data Bank. 3D view of protein was obtained using PyMOL.

## Acknowledgements

The work was funded by grants from the NIH to N.R.L. (DA046100, AI122390 and AI120898) and to M.J.M. (UM1AI148574), T.T. was supported by the Vilcek/Goldfarb Fellowship Endowment Fund.

## Author contributions

T.T. and N.R.L. designed the experiments. H.Z., T.T. and B.M.D. carried out the experiments and analyzed data. T.T., H.Z. and N.R.L. wrote the manuscript. M.I.S. and M.J.M. provided key reagents and useful insights. All authors provided critical comments on manuscript.

## Supplementary figure

**Supplementary Figure 1.**
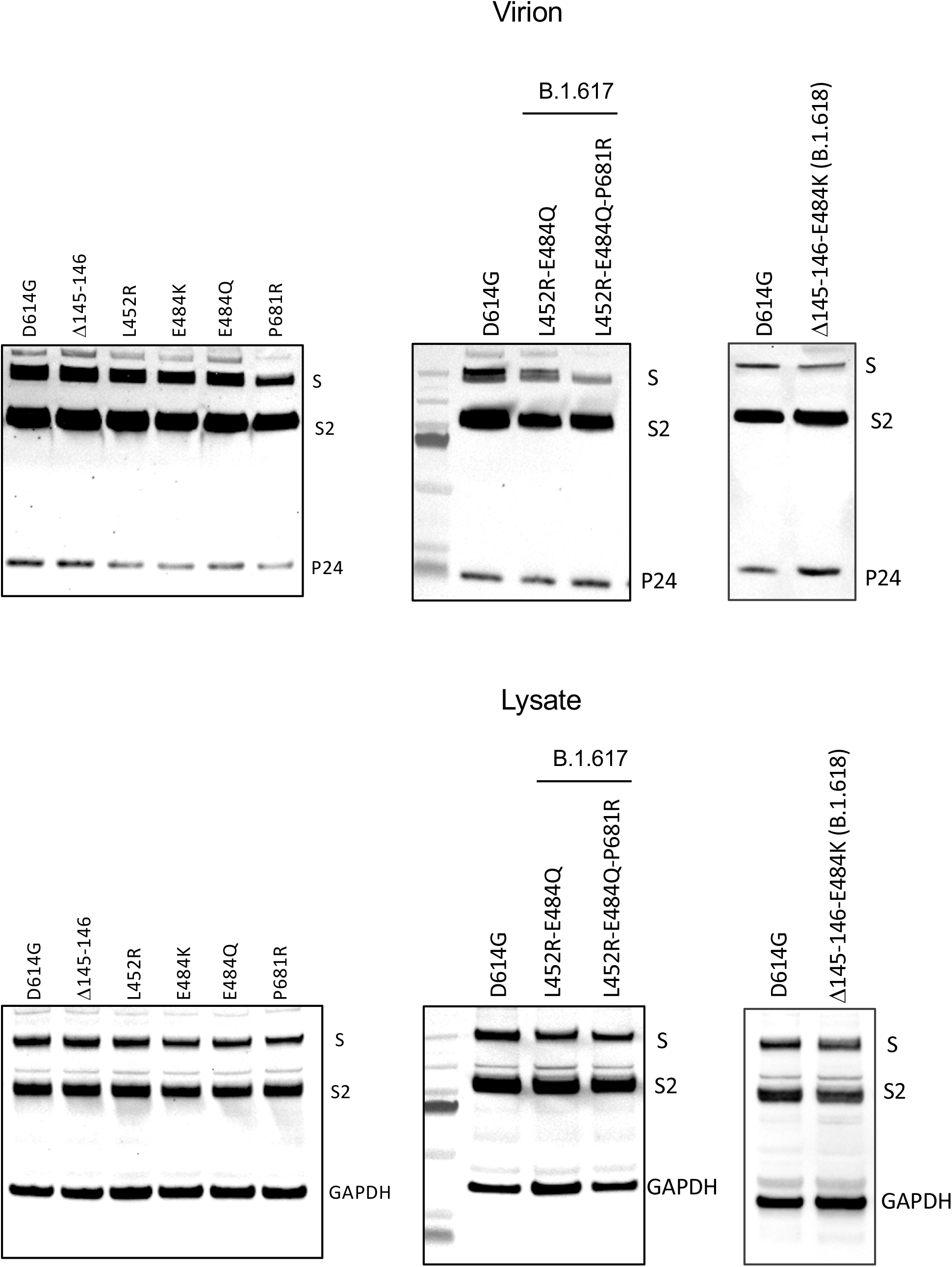
Immunoblot analysis of spike protein in the cellular lysate and lentiviral particles. Pseudotyped viruses were produced by transfection of 293T cells. Two days post-transfection, virions were harvested and analyzed on an immunoblot probed with anti-spike antibody and anti-HIV-1 p24. Cell lysates were probed with anti-spike and anti-GAPDH antibodies.

